# A Glycolysis-Fatty Acid Metabolic Axis Shapes Human Tfh Cell Function

**DOI:** 10.64898/2026.04.01.715763

**Authors:** Someshwar Nath Jha, Bhushan Nikam, Deepak Jena, Shilpa Sachan, Mamuni Swain, Poonam Coshic, Sunil K Raghav, Nimesh Gupta

## Abstract

Follicular helper T (Tfh) cells orchestrate germinal centre-derived humoral immunity by providing essential help to B cells. Despite their key role in humoral immunity, the metabolic processes that guide Tfh differentiation and functions in human remain poorly understood. In this study, we used a human ex vivo Tfh differentiation model to investigate how key metabolic pathways influence Tfh cell differentiation and helper function. Human naïve CD4⁺ T cells were differentiated into Tfh cells, and glycolysis was selectively inhibited during early differentiation to assess its effects on cell fate, function, and transcriptomic landscape. Unexpectedly, early glycolytic inhibition enhanced Tfh cell differentiation but significantly impaired their function, including reduced IL-21 secretion and diminished B cell help. Transcriptomic analyses further revealed downregulation of fatty acid metabolic pathways when glycolysis was inhibited. To better understand this link, we disrupted fatty acid synthesis and oxidation and observed a marked decline in helper functions, including IL-21 and IFN-γ production. Interestingly, acetate supplementation partially restored IFN-γ secretion in glycolysis inhibited conditions, but not IL-21, suggesting that some functional requirements cannot be compensated by alternative metabolic sources. These findings identify glycolysis during early differentiation as a key regulator of human Tfh cells fate and reveal a glycolysis-dependent fatty acid metabolic axis that selectively controls Tfh function. This metabolic checkpoint provides mechanistic insight into tuning Tfh cells and Tfh-driven humoral immunity in vaccination and autoimmunity.

## Introduction

Vaccine-induced protection largely depends on the development of optimal humoral immunity. The breadth and durability of this response are shaped within specialized anatomical structures called germinal centres (GCs). Follicular T helper cells play a crucial role in establishing optimal GC-derived humoral immunity by providing essential help to B cells for their survival, class switching, affinity maturation, and the formation of long-lived memory ^1^. Multiple factors influence Tfh cell development, including the peptide-MHC-TCR dwell time, signal strength, cytokine cues, and transcriptional regulators ^2–4^. While the metabolic axes that govern Tfh cell differentiation and function are poorly understood, emerging evidence suggests that Tfh cells rely on distinct metabolic programs ^5; 6^. As the development of Tfh cells includes their movements in different zones of the secondary lymphoid organ ^7^, it is quite possible that these cells can rely on more than one metabolic pathway to fulfil their biosynthetic demands.

Recent studies using mouse models have revealed a complex and context-dependent role of cellular metabolism in Tfh differentiation and function. mTORC1 signaling exemplifies this duality, while it promotes expression of Blimp-1, a known negative regulator of Tfh lineage commitment, mTORC1 downstream to ICOS is simultaneously required to sustain Tfh function. Similarly, glycolysis appears to play a dual role in Tfh biology. Notably, HIF-1α, a key driver of glycolysis, has been shown to suppress Tfh differentiation, further reinforcing the metabolic complexity of these cells ^8–12^.

Although most of these insights arise from mouse studies, emerging evidence suggests a broader relevance of metabolic pathways in human Tfh-dependent humoral immunity. Leptin can enhance vaccine responses in a Tfh-dependent manner, and individuals with diabetes exhibit disproportionately elevated Tfh cells, pointing to a metabolic link for optimal function of Tfh cells ^13–15^. Collectively, available data indicate that Tfh cells can rely on distinct metabolic programs across contexts. However, it remains unclear how these metabolic circuits are regulating the differentiation and function of Tfh cells. Given the fundamental biological differences between mouse and human in terms of cytokine or signalling cues like neurotransmitter dopamine that govern their differentiation and functions ^16–19^, there is a need to study human Tfh cells independently.

In this study, we investigated metabolic programming in human Tfh cells during different phases of differentiation with a primary focus on glycolysis, which is an essential metabolic switch for T cell activation and has also been linked with BCL6, the master regulator transcription factor for Tfh cells ^20–23^. To dissect the role of glycolysis in Tfh differentiation and function, we used the human *ex vivo* Tfh differentiation model, which successfully mirrors *in vivo* counterparts in terms of the transcriptional signature, producing helper cytokines, and providing help to B cells ^24^. We found that inhibition of glycolysis with 2-deoxyglucose (2-DG) led to increased Tfh differentiation, primarily through enhanced CXCR5 expression. However, this gain came with diminished function including cytokine production and B-cell help. This effect was dominant when glycolysis was inhibited at the initial phase of Tfh differentiation, whereas inhibition of glycolysis later had minimal impact once the Tfh transcriptional program was already established. A detailed transcriptome analyses supported these observations. Glycolysis inhibition enhanced the expression of factors that promote Tfh differentiation, but also increased FOXP1, which may explain the sharp reduction in IL21 production. Interestingly, the transcriptomic signature pointed to a suppressed gene network for fatty acid metabolism. Guided by this clue, we found that glycolysis-inhibited cells have reduced fatty acid metabolism and lower Bodipy uptake. Importantly, blocking fatty acid synthesis or oxidation reduced help function without altering Tfh differentiation, firmly linking fatty acid metabolism to Tfh function. Acetate supplementation partially restored cytokine secretion, further supporting this metabolic axis. Altogether, these findings reveal a metabolic dependency in which human Tfh cells can differentiate under limited glycolysis, but their ability to function requires glycolysis-driven fatty acid metabolism.

## Materials and Methods

### PBMC and Plasma processing

This study was approved by the Institutional Ethical Review Board of the National Institute of Immunology (NII) and the All India Institute of Medical Sciences (AIIMS), New Delhi. We obtained informed consent from all healthy donor volunteers. All the blood samples obtained were processed within 4 hours of collection to isolate PBMCs. Whole blood samples were collected in 350 mL CPD blood bags from all healthy blood donors and then centrifuged at 2000 rpm for 10 minutes to separate the plasma and cellular fractions. Later, 1-2 mL of plasma from the top was carefully removed from the remaining cell fractions and stored in aliquots at −80 °C. PBMCs were separated by Ficoll gradient centrifugation (Ficoll-Paque PLUS, GE Healthcare Life Sciences) and cryopreserved in several aliquots in Fetal Bovine Serum (Gibco) containing 10% (v/v) DMSO (Thermo-Fisher) and stored in liquid nitrogen.

### T cell activation assay

Naïve CD4^+^ T cells were isolated from PBMCs of healthy donor by magnetic bead negative selection with EasySep Naïve CD4^+^ T-Cell Isolation Kit (STEMCELL Technologies). The purity (CD4^+^CD45RA^+^) was 95% or greater. Naïve CD4 T cells (8 × 10^4^ cells/well) were then stimulated with 2µL/well of Dynabeads Human T-Activator CD3/CD28 (Life Technologies) and kept in 5% CO_2_ incubator at 37°C, for the next 5 days. After incubation, cell count and viability was checked using Trypan Blue on LUNA Cell counter

### T-cell Proliferation Assay

The cell proliferation was measured after treatment with 2-DG using CTV (Cell trace violet) dye. 1×10^6^ naive CD4^+^ T cells purified using magnetic sorting (negative selection) were subjected to 5 μM CTV dye (1;1000 dilution, Invitrogen) in PBS and incubated at 37℃ for 20 minutes. After the completion of incubation, cells were washed using 10% FBS RPMI-1640. The cells were cultured with T cell activation bead with the mentioned dose of 2DG and without 2DG as control, and incubated for 5 days, in 5% CO_2_ incubator at 37°C. At the end of the experiment, the proliferation status of the cells was measured on FACS and analyzed using FlowJo. To calculate the proliferation index (PI), the numbers of CTV peaks in each condition were counted (each peak corresponded to 1 generation, the highest intensity peak corresponded to G0 or generation 0, which were undivided cells). The cell number at every generation were also counted, and the Proliferation Index was calculated according to the formula:

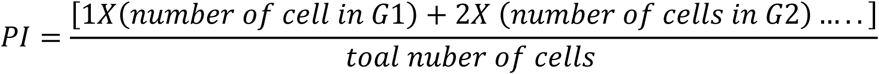

### Ex vivo human Tfh cell differentiation

Naïve CD4^+^ T cells were isolated from healthy donor PBMCs using magnetic bead negative selection with the EasySep Naïve CD4^+^ T-Cell Isolation Kit (STEMCELL Technologies). The purity (CD4^+^CD45RA^+^) was 95% or greater. Naïve CD4 T cells (8 × 10^4^ cells/well) were activated using T cell-Activator beads (CD3/CD28- beads) (2 µL /well, Life Technologies) and cultured with recombinant human activin A (100 ng/ml; R&D) and recombinant human IL-12 (5 ng/ml; R&D) in AIM-V medium (Life Technologies). These cells were then treated with the metabolic pathway inhibitors at the indicated concentrations and kept in 5% CO_2_ incubator at 37°C for the next 5 days. Phenotypes were measured on various days upto 5. Cells were separated using a magnetic separator and finally stained for Tfh-specific markers. Samples were acquired using BD LSRFortessa X-20 and analysed with FlowJo (v.10.7.1).

### Flow Cytometry

The cellular phenotype was examined using Tfh cell specific markers. The CD3/28 activation beads were removed from the culture and cells were washed using 1X PBS before the cells were phenotype at various time of differentiation and in different conditions. Cells were stained for CD4-AlexaFluor 700 (BioLegend), PD-1-PE (BioLegend), CXCR5-BV421 (BioLegend), and Fixable Viability dye-eFlour506 (eBioscience). After adding the antibodies, the cells were incubated in the dark at 4 °C for 1 hour. Cells were washed in FACS buffer and resuspended in 200 μL of buffer before acquiring. Cells were acquired using BD LSRFortessa X-20 and analysed with FlowJo (v.10.7.1).

### ELISpot assays

The IL-21 ELISpot was performed to measure IL21-secreting ex vivo differentiated Tfh cells, with or without treatment of metabolic inhibitors. The ELISpot plate (MSIPS4510; Millipore) was charged with 35% ethanol (Merck Millipore) and coated with the mAb MT216G (10 μg/mL; Mabtech) overnight at 4°C. The plate was washed and blocked with complete RPMI media (having 10% FBS) for 30 min at room temperature. 10,000 cells were seeded along with soluble CD3 and CD28 antibodies (for TCR activation) and incubated in a 5% CO2 incubator at 37 °C for 20 hours. The plate was washed 5 times with PBS and incubated with the mAb MT21.3 m-biotin (0.25 mg/mL; Mabtech) for 2 hours at RT. After incubation, the plate was developed according to the manufacturer’s instructions for the human IL-21 ELISPOT (3540-2A) assay. IL-21 Spots were counted on a vSpot Spectrum ELISPOT reader (AID GmbH) using AID Elispot software v7.0.

### 2-NBDG and Bodipy FL C12 dye uptake assay

The dye uptake assays were performed to determine the kinetics of metabolic pathways during the Tfh cell differentiation in various conditions. The CD3/28 beads were removed fromn the cutured cells using a magnetic bead separator and counted using Trypan Blue and the Luna cell counter. For 2-NBDG staining, 0.2×10^6^ cells were washed and then resuspended in glucose-free media, and 100 μM (final concentration, prepared in glucose-free media) of 2-NBDG dye (Thermo Fisher Scientific) was added to the cell suspension. The exact number of cells was taken and suspended in the AIM-V media for the Bodipy FL C12 (Thermo Fisher Scientific) staining, and 0.1 μM of dye (final concentration, prepared in the AIM-V media) was added to the cell suspension. Both preparations were incubated at 37℃ in CO_2_ incubator for 30 min. Afterwards, the cells were washed with PBS and stained with Tfh-cell specific FACS antibodies: CD4-AF700, PD1-PE, CXCR5-BV421, fixable viability dye eFluor 506. Samples were acquired on a BD LSRFortessa X-20. The flow cytometry results were analyzed using FlowJo software (v.10.7.1).

### Lactate assay

Lactate levels in the cell culture supernatant were measured after the Tfh cell differentiation with or without 2DG. Glycolysis cell-based assay kit was used to detect lactate (Cayman; #600450). Appropriate Lactate standards and controls were used according to the kit instructions. Briefly, 10µL of the supernatant from each condition was transferred to their respective wells, and 100µL of the reaction mixture was added to each well. The reaction mixture includes an appropriate amount of glycolysis substrate, glycolysis cofactor, and enzyme mixture in the provided buffer. This plate was incubated at room temperature with continuous gentle shaking for 30 min. Once the colour developed, absorbance was read at 490 nm using a plate reader. The lactate levels were measured according to the formula provided in the Kit protocol.

### Autologous T-B co-culture

We adapted the previously reported T-B coculture assay protocol to determine the B cell help potential of ex vivo differentiated Tfh cells ^24; 25^. Tfh cell differentiation cultures were set as mentioned above, with or without 2DG. On the 3rd day of culture initiation, autologous memory B cells were isolated from PBMCs (Live CD20^+^CD27^+^). For cell sorting, cryopreserved PBMCs were revived by thawing in a water bath at 37℃ and washed with plain RPMI. After dislodging the pellet, the revived cells were treated with 100 μL/mL of DNase I for 15 minutes at 37 °C. The DNase-treated cells were washed and resuspended in RPMI medium and then counted using Trypan blue. The cells with viability more than 85% were processed further. The resuspended cells were subjected to centrifugation at 1500 rpm for 5 min. At the end of centrifugation, the supernatant was discarded, and antibody cocktails were added for memory B cell sorting: fixable viability dye eFluor 506 (eBioscience), CD20-PE-Cy7 (BioLegend), CD27-PE (BD Bioscience). Cells were incubated in the dark at 4℃ for 1 hour followed by washing with a sorting buffer (1X PBS, 1mM EDTA, and 2% FBS). Cell pellets were resuspended in 3 mL of sorting buffer and filtered with a 40 µM cell strainer. Cells were sorted on BD FACS Aria Fusion. Meanwhile, beads were removed from the Tfh culture, and cells were counted. The sorted memory B cells were co-cultured with the differentiated Tfh cells in a fixed ratio of 100:1. For T cell-B cell collaboration, 0.25 ng/mL of SEB was added to the final volume of 200 μL and kept in the CO_2_ incubator for the next 7 days. On the 7th day, the culture was stained for CD20-PE-Cy7 (BioLegend), CD27-PE (BD Biosciences), CD19-BV785 (BioLegend), CD38 (PE-Cy5, BD Bioscience) and Fixable viability dye eFlour 506. On the 7^th^ day of the coculture, plasma cells (CD19^+^CD20^low^ CD38^high^) were measured using flow cytometer, and co-culture supernatants were stored at - 20℃ for measuring IgG titers.

### ELISA

The antibody titers in autologous co-cultures were measured using the sandwich ELISA. Goat anti-human Fab antibody (SouthernBiotech) was coated on an ELISA plate at a concentration of 0.5 µg/mL (50 µL per well, in 1X PBS) and incubated overnight at 4℃. The plate was washed and blocked using 200 μL of blocking buffer (3% skimmed milk in 0.05% PBST) to each well and incubated for 2h in the dark at room temperature. Meanwhile, culture supernatants are diluted to the desired concentration (here 1:30). After blocking, washing was performed with 200 µL wash buffer and diluted samples were added to the designated wells. The plate with the sample was incubated for 1.5 h in the dark, and after the incubation time, the wells were again washed as in the previous step. 100 μl of HRP-conjugated anti-human Fc-IgG (1:4000; SouthernBiotech) was added to each well and incubated in the dark for 1h at room temperature. After incubation, wells were washed, and 100 μL of substrate (OPD, Sigma-Aldrich) was added to each well. Following the colour development 2N HCl was added to stop the reaction. Absorbance was taken at 492nm on Multiskan Go plate reader (Thermo Scientific).

### Acetate Supplementation Assay

Human naïve CD4^+^ T were cultured with Tfh differentiation milieu as stated above in presence or not of 2DG. 2DG-treated wells were supplemented with 5 mM of acetate, and at the end of the experiment, cells were examined for Tfh differentiation and function. The Tfh functions were determined as their ability in producing IFN*γ* and IL-21 by intracellular cytokine staining (ICS) assay and ELISpot, respectively.

### Intracellular Cytokine Staining (ICS) Assay

Cells were stimulated as per the experimental requirements, and at the end of experimental time-points, cells were treated with PMA (25ng/mL, Sigma Aldrich) and ionomycin (1 μg/mL, Sigma Aldrich) along with GolgiPlug containing Brefeldin A (0.5 μL/well, BD Biosciences) for 4 hours. After the end of 4 hours, cells were first stained with surface markers antibodies as mentioned in Flow cytometry section and followed by this, cells were fixed and permeabilized for 30 min and washed with 1 mL permeabilization buffer (FOXP3/Transcription factor buffer Kit, Thermo Fisher Scientific). After fixation and permeabilization, antibody mix prepared in PERM buffer for cytokines were added (IFN*γ*; PE-CF594 and IL21; AlexaFlour 647; BD Bioscience) and incubated at 4℃ in dark for 1 hour. The cells were finally washed with the PERM buffer and resuspended in 200 μL of FACS buffer. All the sample were acquired on BD LSR Fortessa X-20 and were analyzed using FlowJo software (v.10.7.1).

### Transcriptomic analysis

For RNAseq library preparation, high-quality total RNA was first extracted from the Tfh cells under two different conditions of with or without 2-DG treatment and quantified to ensure RNA integrity and concentration. mRNA was isolated from 500ng of total RNA using mRNA isolation module (mRNA isolation module, NEB), which selectively captures polyadenylated transcripts using oligo-dT-conjugated magnetic beads. mRNA library preparation was carried out using the NEB Next Ultra II RNA library preparation kit according to instructions manual. The isolated mRNA was fragmented with a target of 200nt and primed for first strand cDNA synthesis. Random primers and reverse transcriptase were used for the synthesis of the first-strand cDNA. Second-strand synthesis was conducted using a combination of DNA Polymerase I and RNase H. The double-stranded cDNA underwent end repair to generate blunt ends. dA-tailing was done to facilitate adapter ligation and then indexed adapters (Illumina-compatible) were ligated to the cDNA fragments. Ligation products were size-selected and purified using SPRIselect beads, which remove small fragments and adapter dimers. The adapter-ligated cDNA fragments were enriched using PCR, followed by a final bead-based purification. After amplification, the concentration of libraries was checked using Qubit 4.0 (Invitrogen) and fragment sizes were analysed on Bio-analyser (Agilent). cDNA libraries were denatured using 0.2N NaOH at RT. Further, Tris-Cl pH 8.5 was used to neutralise the effect of NaOH. cDNA libraries were diluted using HT1 buffer to a final concentration of 1.7pM, which were used for loading. Subsequently, libraries were sequenced by Illumina NextSeq 550 platform.

### RNAseq data analyses

Quality of the paired-end raw reads was checked using FastQC tool, and then aligned to human reference genome (hg38) downloaded from the UCSC genome browser using STAR^26^. Gene counts were extracted from the aligned bam file using featureCounts function from SubRead package ^27^. Further, Principal component analysis and differential gene expression analysis were performed using DESeq2 package ^28^. ComplexHeatmap package was used to plot heatmap of the expression of target genes ^29^. To compare normalized count between two groups ggplot2 and ggpubr (v 0.6.0) R packages were used. Paired T-test statistics was used for comparison between two groups.

### Qualitative real-time PCR

The differentiated Tfh cells with or without 2-DG treatment were collected on day 5. Cells were washed with PBS and resuspended in RLT plus buffer. RNA was extracted using RNAeasy Plus Micro Kit (Qiagen) according to the kit protocol. The RNA was dissolved in 20µl RNase-free DEPC water. RNA was quantified using a Multiskan Sky High microplate spectrophotometer (Thermo Fisher), and a 260/280 ratio was used to determine RNA quality. The cDNA was then synthesised from 1μg of total RNA using the Reverse Transcription Kit (Takara) according to the manufacturer’s protocol. Each sample was amplified in triplicate with primers specific to genes of interest using 15 ng of cDNA. A Quant Studio 4 real-time PCR system was used to detect gene expression. Results were normalised to 18s gene expression, and the expression of the target gene was calculated as the “fold change” relative to the control samples according to the 2^−ΔΔ*C*t^ method using the formula:

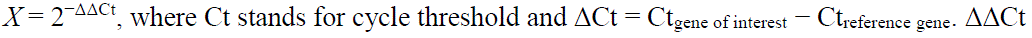

is the difference between the ΔCt values of the “experimental” samples and the ΔCt of the corresponding “vehicle/control” sample.

Primer sequences: BCL6- Forward: CATGCAGAGATGTGCCTCCACA; Reverse: TCAGAGAAGCGGCAGTCACACT. PRDM1/BLIMP1- Forward: CAGTTCCTAAGAACGCCAACAGG; Reverse: GTGCTGGATTCACATAGCGCATC. HK2- Forward: GAGTTTGACCTGGATGTGGTTGC; Reverse: CCTCCATGTAGCAGGCATTGCT. IL-21- Forward: CCAAGGTCAAGATCGCCACATG; Reverse: TGGAGCTGGCAGAAATTCAGGG. CXCR5- Forward: TGAAGTTCCGCAGTGACCTGTC; Reverse: GAGGTGGCATTCTCTGACTCAG. 18s- Forward: ACCCGTTGAACCCCATTCGTGA; Reverse: GCCTCACTAAACCATCCAATCGG.

## Results

### Glycolysis inhibition enhances Tfh differentiation but suppresses its function

To investigate the metabolic requirement for differentiation and function, we utilised the ex vivo human Tfh differentiation model ^24^. We first examined the role of glycolysis in Tfh differentiation by perturbing glycolysis using a glucose analogue 2-deoxyglucose (2-DG). The working concentration of 2DG was determined by activating human naïve CD4^+^ T cells using CD3 and CD28 activation beads, with or without various concentrations of 2DG from 1 to 4 mM. 2DG was found to be toxic for activated T cells at higher concentrations (Figure 1A). The effectiveness of 2DG was further validated in a T-cell proliferation assay, and a lower concentration of 1 mM was selected for further investigations (Figure 1B). The significant decline in the lactate levels, by-product of glycolysis, in cell supernatant from 2DG treated conditions confirm the effectiveness of glycolysis inhibition at the selected dose (Figure S1B). We then examined the effect of glycolysis inhibition on the human Tfh cells differentiated from the naïve CD4^+^ T cells in Activin A and IL12 supplemented conditions, along with CD3-CD28 activation beads for 5 days (Figure S1A). The glycolysis-inhibition led to a significant increase in the Tfh cells differentiation as compared to glycolysis-sufficient condition (Figure 1C). The expansion of Tfh numbers was associated with the increased expression of CXCR5 (Figure 1D) but not PD1 (Figure 1E). The significantly increased frequency of Tfh cells in the presence of Glut inhibitor (Glutor) further confirmed the positive impact of inhibited glucose uptake and glycolysis-inhibition on Tfh-cell differentiation (Figure S1C).

**Figure 1:**
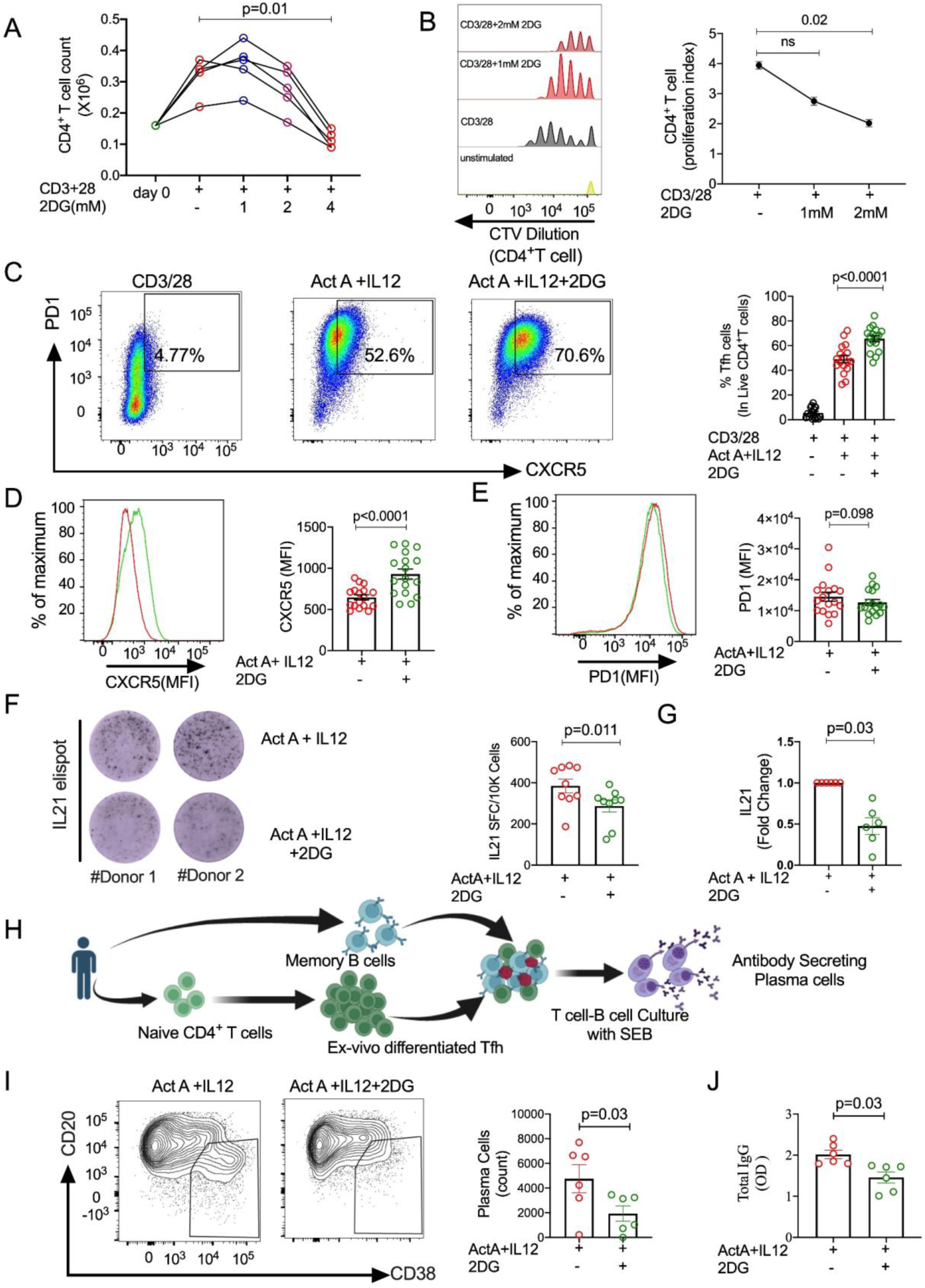
Inhibition of glycolysis leads to enhanced Tfh cell differentiation but compromised function. **(A)** Dose standardisation of 2DG with the activated CD4^+^ T cells. Viable cell counts were plotted after day 5 of activation in the presence of 2DG (n=5). Each line represents one individual. **(B)** Inhibition of proliferation of CD4^+^ T cells after 2DG treatment (n=3) in a CTV dye assay. **(C)** Representative FACS plot alongside percentage of follicular T helper cells in live CD4^+^ T cells differentiated with Tfh differentiation milieu with or without glycolysis inhibition (n=17). **(D-E)** Mean Fluorescence intensity (MFI) of CXCR5 and PD1 in Tfh cells differentiated with or without glycolysis inhibition (n=17). **(F)** Frequency of IL-21 producing Tfh cells, cultured in Tfh differentiation milieu with or without glycolysis inhibition (n=9). **(G)** IL-21 RNA levels measured by qPCR in cells cultured in Tfh differentiation milieu with or without glycolysis inhibition (n=6). **(H)** Scheme depicting the autologous Tfh-B cell co-culture assay. **(I)** Number of plasma cells in autologous Tfh-B cell co-cultures, with Tfh cells differentiated with or without glycolysis inhibition (n=6). **(J)** IgG level as measured by ELISA in the culture supernatant of the above stated Tfh-B cell co-culture (n=6). Data represented as mean±SEM. Statistics: **(A-B)** Friedman Test followed by Dunn’s correction, **(C-J)** Wilcoxon matched pair signed rank test.

IL-21 is the major helper cytokine secreted by Tfh cells to exert its help to B cells. Therefore, to check whether the increase in Tfh differentiation also implies the increase in functional potential of these cells, we probed the level of IL-21 secreted by Tfh cells with or without glycolysis-inhibition. Surprisingly, glycolysis-inhibition led to a significant decline in IL-21 secreting cells, which was evident both at protein (Figure 1F) and transcript levels (Figure 1G). Intrigued by these findings, we then performed an autologous Tfh-B co-culture assay to determine the B cell help potential of Tfh cells. The ex vivo differentiated Tfh cells with or without glycolysis inhibition were collected at Day 3 of culture, washed and co-cultured with autologous memory B cells, followed by stimulation with SEB. The plasma cell differentiation and the antibody levels were measured as the outcome of Tfh help potential to memory B cells (Figure 1H). The Tfh cells from glycolysis-inhibited 2DG treated condition showed compromised help potential as marked by significant decline in plasma cell differentiation (Figure 1I) as well the reduced IgG levels in the co-culture supernatant (Figure 1J). Altogether, these findings suggest that while glycolysis inhibition may promote Tfh cell differentiation, it compromises their hallmark function of providing help to B cells.

### Glycolysis-inhibition during the early phase of acquiring Tfh program is essential for enhanced Tfh differentiation and suppressed function

We next investigated the kinetics of Tfh cells differentiation to determine the impact of glycolysis-inhibition from Day 1 to Day 5 (Figure 2A). Interestingly, glycolysis-inhibition led to a suppression in Tfh cells frequency at Day 2 (Figure 2B), which was also evident by the decline in expression levels of CXCR5 (Figure 2C) and PD1 (Figure 2D). However, this suppression was only evident until day 2, beyond which the frequency of Tfh cells significantly increased Day 3 onwards in the glycolysis-inhibition conditions over the control (Figure 2B). A similar trend was found with the increase in expression of CXCR5 (Figure 2C) and PD1 (Figure 2D). We next investigated the effect of glycolysis-inhibition at different phases of Tfh differentiation. The glycolysis was inhibited either at the initial phase of day 0 or during the later phase of Tfh differentiation, at Day 3 (Figure 2E). The inhibition of glycolysis at initial phase (Day 0) resulted in enhanced Tfh frequency as seen previously, but not the treatment at Day 3 (Figure 2F). In fact, the 2DG treatment at Day 3 showed no significant effect on CXCR5 and PD1 expression (Figure 2G-H). Moreover, the Tfh function assessed by measuring IL-21 levels revealed no significant effect when the glycolysis was inhibited at the later phases of Day 3 (Figure 2I).

**Figure 2:**
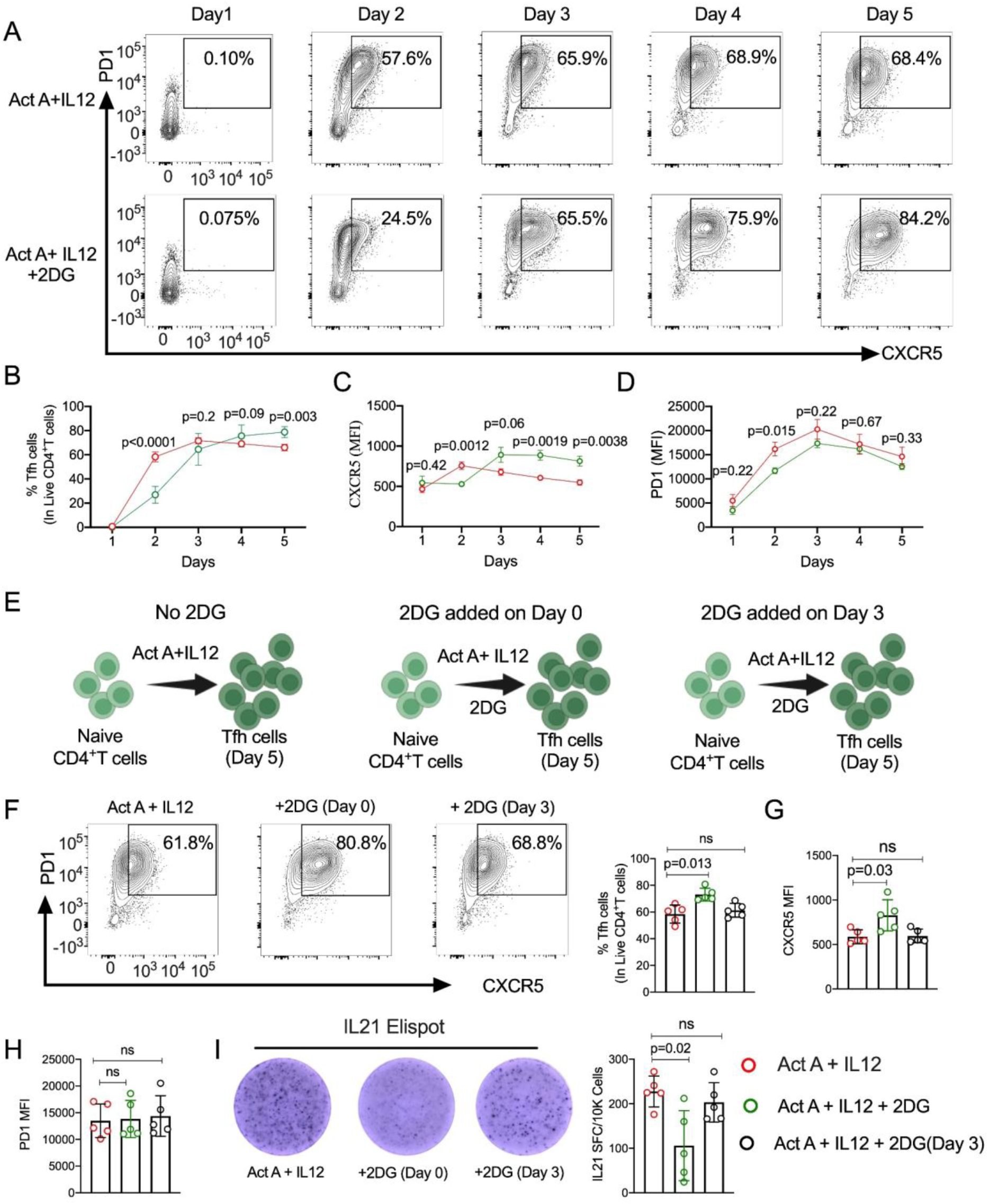
Early glycolysis inhibition is essential for impacting Tfh cell differentiation and function. **(A)** Representative FACS plot of Tfh cells analyzed at indicated time points in glycolysis inhibited condition. **(B)** Percentage of Tfh cells in the Live CD4^+^T cells differentiated under Tfh-differentiated milieu with or without glycolysis inhibition (n=6) **(C-D)** MFI of CXCR5 and PD1 in the Tfh cells at the different time points as in (B) (n=6). **(E)** Experimental setup to investigate the effect of glycolysis inhibition at different time points during Tfh differentiation. **(F)** Percentage of Tfh cells in live CD4^+^T cells cultured as in (E) (n=5). **(G-H)** MFI of CXCR5 and PD1 in Tfh cells (n=5). **(I)** Frequency of IL-21 producing Tfh cells, cultured in Tfh differentiation milieu as in (E) (n=5). Data sets are presented as mean ±SEM. Statistics: **(B-D)** multiple T-tests. **(E-I)** Friedman test, followed by Dunn’s correction.

These results suggest that early inhibition of glycolysis is essential for an increased Tfh differentiation in the later time points, despite of a transient suppression at initial stage. Notably, early metabolic disruption causes lasting damage to their ability to function effectively. Interestingly, glycolysis-inhibition has no impact on differentiation or function after the Tfh program has already been established.

### Inhibition of glycolysis using 2DG lead to a significant change in the Tfh transcriptomic program and perturbation of fatty acid metabolism

Next, to examine how glycolysis inhibition impacts the Tfh-cell program and its functions, we performed transcriptomic analyses under conditions with and without glycolysis inhibition. The PC analysis of the sequenced library showed a clear segregation between the RNA of the CD4^+^ T cells differentiated to Tfh with or without 2DG (Figure 3A). We did differential expression gene analysis of 5 donors and found significant number of differentially regulated genes after 2DG treatment (Figure 3B; Supplementary Table S1). There was marginal donor-to-donor variation in the expression pattern of the genes (Figure 3B). Therefore, we did pair-wise analysis to uncover the change in expression of genes involved in Tfh differentiation, function, and localization. Here, we found that glycolysis inhibition using 2DG showed promoting effects on the Tfh differentiation, which corroborated with our above observations. We found that genes that negatively regulate the differentiation of Tfh are downregulated and the genes that support the Tfh differentiation were upregulated after 2DG treatment (Figure 3C). Negative regulators like PRDM1, KLF2, and IL2RB were downregulated after glycolysis-inhibition (Figure 3C & E). We observed a similar trend in BCL6 and CXCR5, where 2DG treatment also led to enhanced levels of these Tfh-defining factors compared to non-treated conditions (Figure 3C). Interestingly, the Tfh program specific coordinated transcriptional signature was evident. For instance, ASCL2, which is critical for downregulating PRDM1, KLF2, and S1PR1 and thus supports GC-Tfh differentiation ^30^, was found to be upregulated in the glycolysis-inhibited conditions as compared to the non 2DG-treated cells (Figure 3C & E). It is worth noting that ASCL2 is required for upregulating ITCH, and ITCH, in turn, downregulates FOXO1 further to support the Tfh differentiation ^30; 31^. We also observed a similar trend, where 2DG treatment led to the upregulation of ITCH and the downregulation of FOXO1 (Figure 3C).

**Figure 3:**
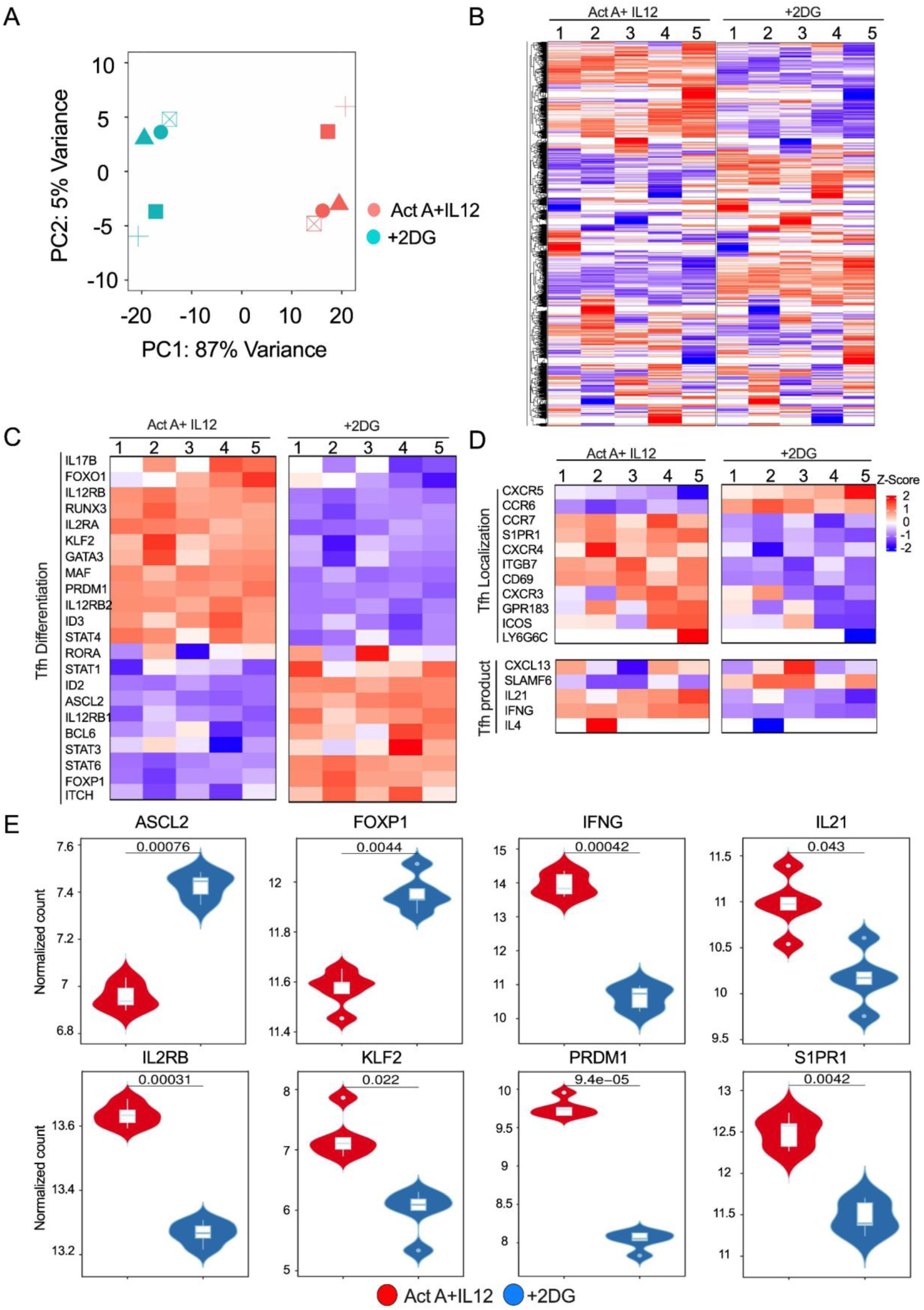
Glycolysis inhibition induces significant alterations in the Tfh cell program and transcriptional landscape. **(A)** PC analysis summarizing the difference between transcriptomes of CD4^+^ T cells cultured in Tfh differentiation milieu with or without glycolysis inhibition. **(B)** Heatmap showing the overall transcriptional profile of CD4^+^ T cells cultured as in (A). **(C)** Heatmap showing differential expression of Tfh-specific genes in the two above conditions. **(D)** Heatmap, showing differential expression of Tfh localization and function-specific genes. **(E)** Graph of some of the highly affected genes responsible for Tfh localization, differentiation, and functions. Complex heatmap function is used to plot heatmaps, Data set is represented as mean ±SD, n=5. Statistics: Pair T-test.

Although inhibition of glycolysis promoted Tfh differentiation, FOXP1 levels in cells treated with 2DG were higher compared to the non-treated cells (Figure 3C & E). Interestingly, FOXP1 plays a critical role in suppressing Tfh cytokines like IL-21 and IFNγ ^32–34^ and therefore, it’s lower expression is evident with potent functional Tfh cells.

Another notable feature of Tfh differentiation under glycolysis-inhibited conditions was the coordinated modulation of genes that regulate the spatial localization of Tfh cells within germinal centers. Transcriptomic profiling revealed a distinct enhancement of the CXCR5-axis, which is essential for the migration of Tfh cells toward B cells follicles. Consistent with this, we observed a concomitant downregulation of CCR7, a prerequisite for efficient CXCR5-mediated positing of Tfh cells (Figure 3D). Strikingly, glycolysis inhibition also resulted in reduced expression of S1PR1 together with its upstream transcriptional regulator KLF2, a pattern strongly indicative of activation of the ASCL2 regulatory program, that enforces follicular commitment and GC localization^30; 35^.

Furthermore, inhibition of glycolysis led to a downregulation of CD69 and an upregulation of CCR6 (Figure 3D). CD69 expression promotes retention of Tfh cells within secondary lymphoid organs and suppresses chemotactic responsiveness through S1PR1 downregulation ^36^. Notably, CD69 is metabolically regulated as a direct target of HIF-1alpha and reduced glycolysis likely contributed to its downregulation ^37^. Corroborating with the above results on Tfh functionality, we also found a significant downregulation of IL-21 and IFNγ transcripts (Figure 3D & E).

Taken together, these findings reveal that perturbing glycolysis reinforces the ASCL2-CXCR5 axis leading to increased Tfh cell differentiation and enhanced migratory potential towards B-cell follicle, while their functional ability was compromised.

### Inhibition of glycolysis resulted in perturbation of fatty acid metabolism in Tfh cells

As the Tfh cells showed enhanced differentiation despite of glycolysis-inhibition, we examined the transcriptomic profiles of metabolic pathways including glycolysis. Some of the genes from the glycolysis pathway showed increased expression including HK1, HK2, PFKL, LDHB, ALDOA (Figure 4 A & B). Even targeted qPCR showed similar expression pattern of these genes including HK2 (Supplementary Figure 3D). Likewise, a similar trend was observed in case of the genes involved in TCA cycle like MDH1, PDHA1 (Figure 4C & D).

**Figure 4:**
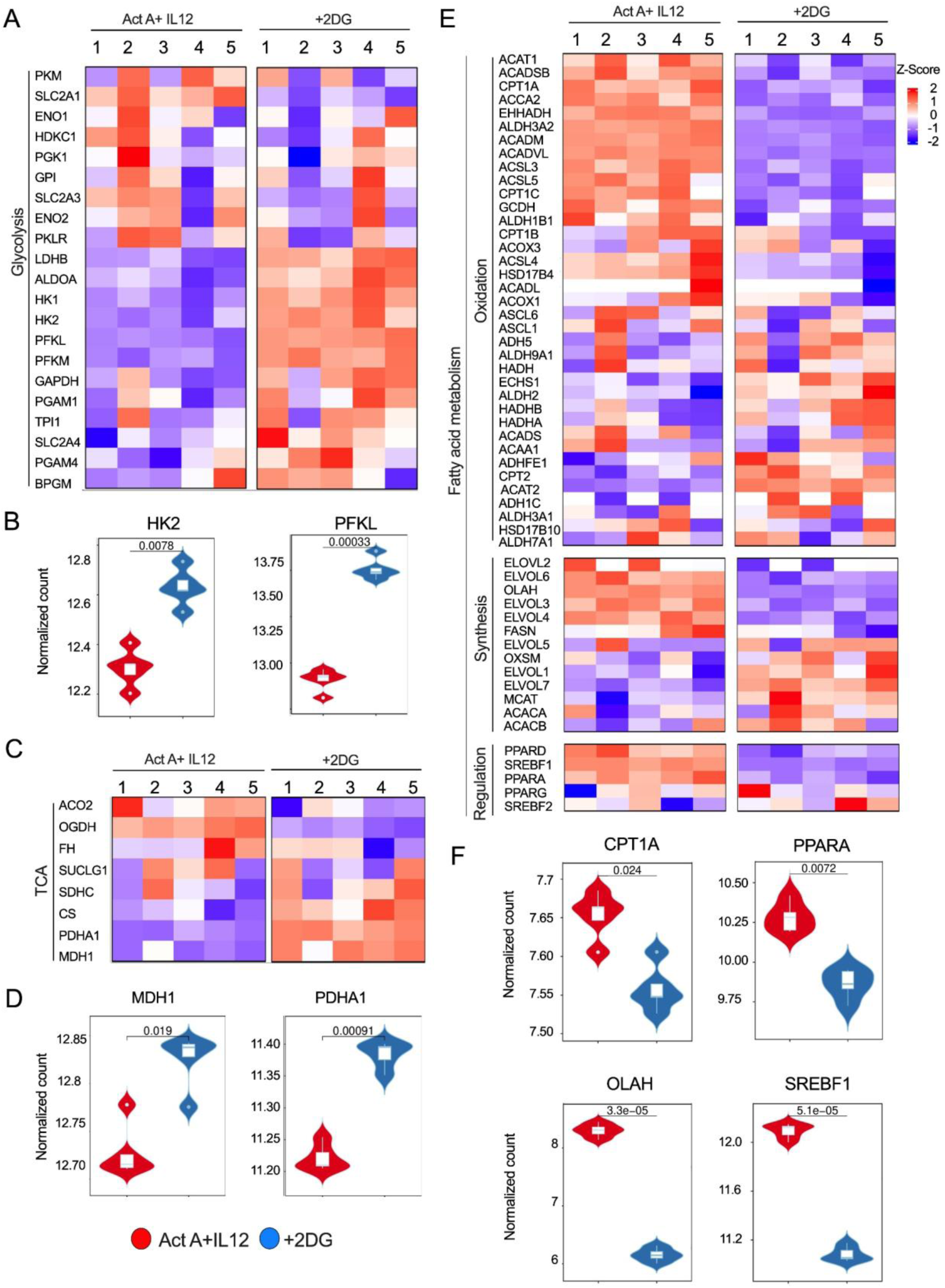
Glycolysis inhibition triggers metabolic reprogramming in Tfh cells. **(A-B)** The heatmap shows differentially expressed genes related to glycolysis in Tfh cells with or without glycolysis inhibition, and graphs of selected genes showing change in expression level. **(C-D)** Heatmap of genes showing differentially expressed genes of the TCA cycle after 2DG treatment and representative graphs of selected genes in the same condition as in (A). **(E-F)** Heatmap showing differential expression of genes related to Fatty acid metabolism in Tfh cells with or without glycolysis inhibition. Data set is represented as mean ±SD, n=5. Statistics: Pair T test.

As glucose can be utilized for de novo fatty acid synthesis, inhibition of glycolysis can severely impact fatty acid metabolism. Glycolysis-dependent lipogenesis was pivotal in determining the fate of Th17 cells ^38^. Therefore, we examined at the impact of inhibition of glycolysis on genes that are involved in fatty acid metabolism. Surprisingly, a severe negative impact of glycolysis inhibition was found on the transcriptomic landscape that is involved in fatty acid metabolism. This impact was observed on the genes that are crucial for fatty acid oxidation (CPT1), elongation (OLAH), and for the regulation of fatty acid metabolism (PPARA and SREBF) (Figure 4E), with the strongest impact on OLAH (Figure 4E & F).

These results suggested a possibility of a feedback response from the cells to compensate the decline in productive glycolytic output. Importantly, the glycolysis mediated regulation of fatty acid metabolism was evident, as the glycolysis-inhibition led to the significant downregulation of gene network for fatty acid metabolism.

### Fatty acid metabolism regulates Tfh cells functionality

Transcriptomic analyses revealed a significant disruption of fatty acid metabolic pathways following glycolysis-inhibition. We therefore further examined the fatty acid metabolism in the Tfh cells. Tfh cells showed a significantly higher uptake of Bodipy (Fl-C12 fatty acid) as compared to the non-Tfh cells (PD1^-^CXCR5^-^), indicating a high dependability on fatty acid metabolism during the differentiation (Figure 5A; Supplementary Figure 4A). Tfh cells showed a significant decline in the fatty acid levels in glycolysis-inhibited conditions (Figure 5B), which was not the case with a glucose analogue (2-NBDG) uptake by these cells (Supplementary Figure 4B-C). The kinetics analyses revealed a higher fatty acid level throughout the Tfh differentiation cycle from Day 2 to Day 5 as compared to the Tfh differentiation in glycolysis-inhibition condition (Figure 5C).

**Figure 5:**
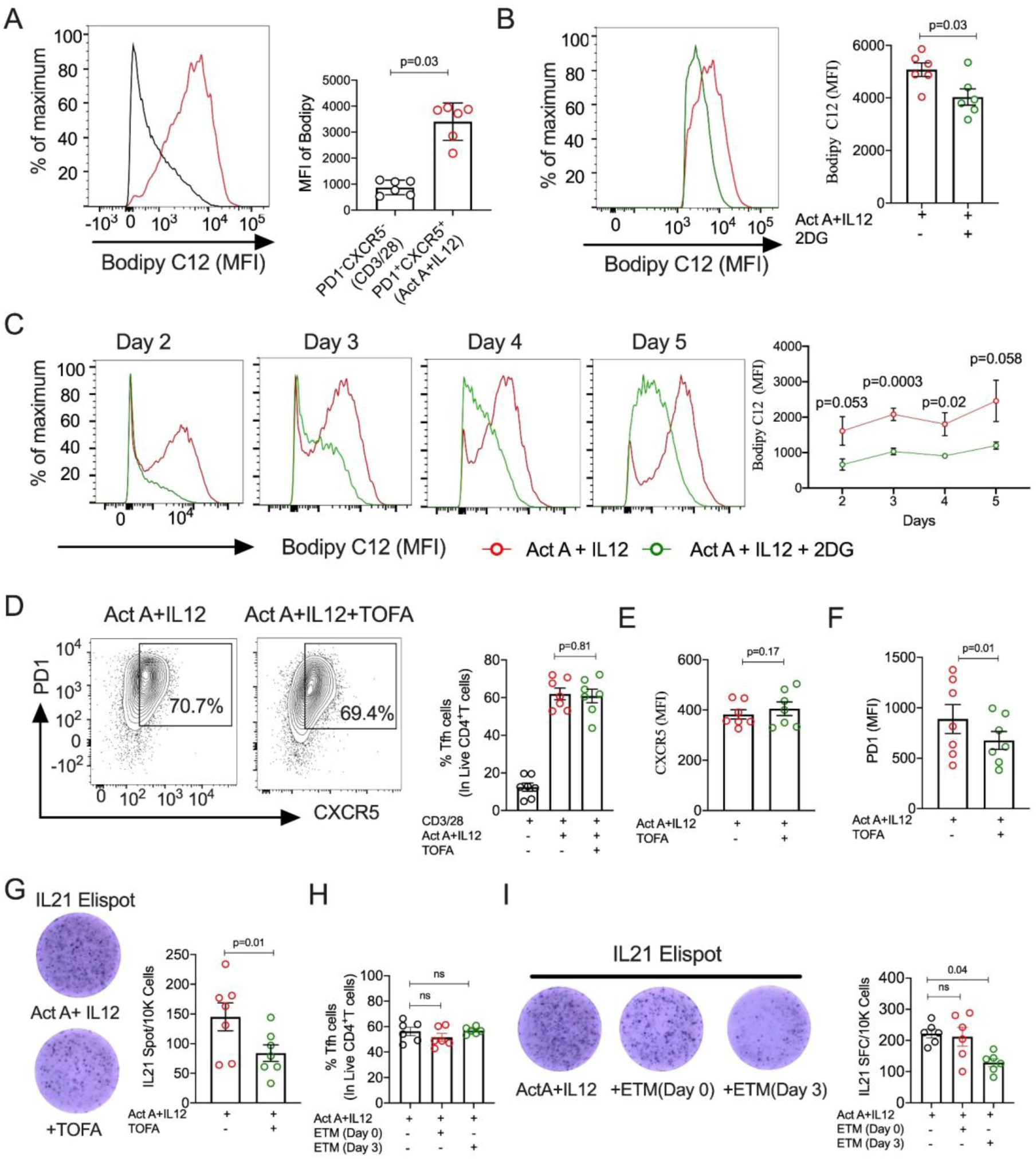
Inhibition of Fatty Acid Synthesis and Oxidation impairs Tfh cell function. **(A)** Difference in MFI of Bodipy FL-C12 in Tfh cells, compared to non-Tfh (PD1^-^CXCR5^-^). **(B)** Bodipy FL-C12 MFI i Tfh cells, differentiated with or without glycolysis inhibition. **(C)** Bodipy FL-C12 level in Tfh cells at different time points when cultured with or without glycolysis inhibition. **(D)** Effect of TOFA (fatty acid synthesis inhibition) on Tfh differentiation. **(E-F)** Change in MFI of CXCR5 and PD1 in Tfh cells after inhibiting fatty acid synthesis. **(G)** IL-21 Spot formed by CD4^+^T cells cultured in Tfh differentiation milieu, along with fatty acid synthesis inhibition. **(H)** Effect of inhibition of fatty acid oxidation at indicated time points using Etomoxir on Tfh differentiation. **(I)** IL-21 Spot formed by CD4^+^T cells cultured in Tfh differentiation milieu, along with fatty acid oxidation inhibition at indicated time points. Data sets are presented as Mean ± SEM, n=6 in each set of experiments. Statistics: (A-B, D-F), Wilcoxon matched-pair signed rank test. (C) multiple T-test. (H-I) Friedman test, followed by Dunn’s correction.

As the above data supported significant perturbation of the genes involved in fatty acid synthesis, we next inhibited fatty acid synthesis using TOFA (5-(Tetradecycloxy)-2-furoic acid), a competitive and reversible inhibitor for Acetyl-CoA carboxylase. The naïve CD4^+^ T cells were cultured with Tfh differentiation milieu with or without TOFA (1μM). Inhibition of fatty acid synthesis showed no significant effect on Tfh differentiation (Figure 5D). While a marginal decline was observed in the PD1 expression, the CXCR5 expression remained unaffected (Figure 5E & F). Interestingly, inhibition of fatty acid synthesis alone showed a significant decline in the Tfh functions, as measured by the levels of IL-21 (Figure 5G). We further examined the effect of fatty acid oxidation on Tfh cells differentiation by blocking FAO using Etomoxir. Similar to fatty acid synthesis, inhibition of fatty acid oxidation showed no significant impact on Tfh cell differentiation (Figure 5H). However, when Etomoxir was added on day 3, but not at the early phase on day 0, a marked reduction in IL-21 production by Tfh cells was observed (Figure 5I).

Altogether, these findings highlight the dependence of Tfh cells on fatty acid metabolism, which can be disrupted by glycolysis inhibition. Notably, fatty acid metabolism showed no implication with the Tfh cell differentiation, but its critical role in regulating Tfh cell function was clearly evident.

### Acetate supplementation partially restores Tfh function in glycolysis-inhibited conditions

We hypothesized that 2-DG treatment led to reduced levels of acetyl-CoA, a key glycolytic intermediate essential for de novo fatty acid synthesis. Acetyl-CoA is carboxylated by acetyl-CoA carboxylase (ACC) to form malonyl-CoA, which is a rate-limiting step in fatty acid biosynthesis ^39; 40^. We thus evaluated the effect of acetate supplementation on Tfh cells under glycolysis-inhibited conditions. Acetate supplementation showed no significant effect on Tfh cell differentiation compared to cells with glycolysis inhibition alone (Figure 6A). Interestingly, it partially restored IFNγ secretion during the glycolysis inhibition (Figure 6B). However, acetate supplementation during glycolysis-inhibited condition failed to rescue IL-21 production, as assessed by both ICS and ELISpot assay (Figure 6C-E).

**Figure 6:**
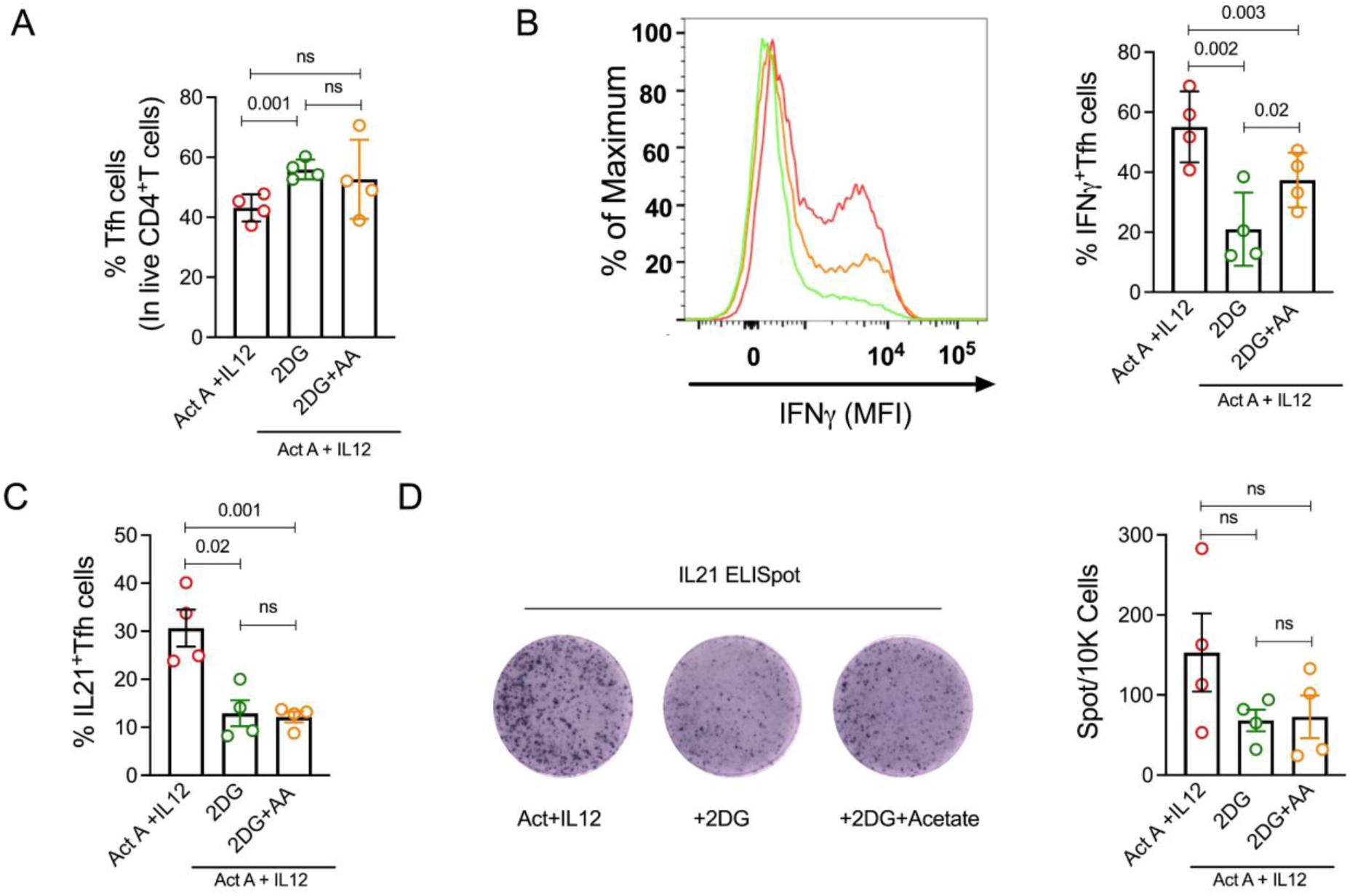
Acetate supplementation partially restores Tfh cell function. **(A)** Frequency of Tfh cells in glycolysis-sufficient, glycolysis-inhibited, and acetate-supplemented glycolysis-inhibited Tfh cell differentiation culture. **(B-C)** Comparison of IFNγ and IL21 levels as detected through intracellular staining in Tfh cells with or without acetate supplementation in glycolysis-inhibited conditions. **(D)** Frequency of IL-21 producing Tfh cells, along with glycolysis inhibition supplemented with acetate. Data are represented as mean±SEM, (n=4). Statistics: (A-D) Friedman test, followed by Dunn’s correction.

Together, these observations suggest that acetate supplementation cannot rescue the ability of Tfh cells to produce IL-21, however, it may partially rescue the function of Tfh cells in producing IFNγ in glycolysis-restricted conditions.

## Discussion

In this study, we have uncovered a unique metabolic signature of the Tfh cells, which is highly dependent on fatty acid metabolism for its function. Interestingly, fatty acid metabolism in these cells depends on glycolysis, and hampering glycolysis profoundly affects both fatty acid metabolism and its function. We further established that the effects of glycolysis inhibition could be rescued partially via supplementation of acetate, leading to an improvement in the ability of Tfh to produce IFN*γ*, but not IL-21.

Surprisingly, glycolysis inhibition led to a transient suppression of Tfh cells in the initial days, followed by an expansion of these cells at the later stages. While these cells gained an increase in differentiation, their functional capability was absolutely suppressed. The increase in Tfh differentiation can be explained by upregulated BCL6 and downregulated BLIMP1 (PRDM1), the axis that drives the acquisition of Tfh program ^2^.

Other transcriptional axes that were affected by glycolysis inhibition include ASCL2 and KLF2. ASCL2 can promote CXCR5 expression independent of BCL6 and suppress CCR7. Both of these factors are critical for the migration of Tfh toward the germinal centre. Furthermore, ID3 can negatively regulate ASCL2, indicating that the upregulation of ASCL2 must coincide with downregulated ID3 and CCR7, but upregulated CXCR5 ^30^. The similar axis was found active in the glycolysis-inhibited Tfh cells. It’s possible that the strong induction of this axis supported the enhanced Tfh differentiation observed in glycolysis-inhibited conditions. In contrast to ASCL2, KLF2 is a proven suppressor of Tfh cell transcriptional program, and it does so by upregulating BLIMP1 and S1PR1^35^. The negative effect of KLF2 on Tfh is not only limited to this, as it can also promote non-Tfh fate by upregulating GATA3 ^35^. Thus, the glycolysis-inhibition mediated suppression of KLF2 might have further promoted the Tfh cell differentiation program. Another feature of the glycolysis inhibition that supports enhanced Tfh differentiation is downregulation of FOXO1, a negative regulator of Tfh that can directly bind to the BCL6 gene ^3^ FOXO1 can be negatively regulated by ITCH (an E3 ubiquitin ligase), which promotes FOXO1 protein degradation through ubiquitination ^31^, and thus ITCH-mediated downregulation of FOXO1 can support Tfh differentiation, which was reflected in our transcriptomic analysis.

Hence, besides increased expression of BCL6 and decreased BLIMP1, a downregulated KLF2, and FOXO1, and upregulated ASCL2, and ITCH might be responsible for enhanced Tfh cell differentiation. Interestingly, this observation is accompanied by an upregulation of FOXP1, which may explain the reduced functional potential of these cells. Beside acting as a key negative regulator of Tfh differentiation, FOXP1 can directly bind to the promoter region of IL-21 and suppress its transcription, thereby limiting IL-21 production by Tfh cells ^32; 34^. FOXP1 may similarly influence IFN*γ* secretion in Tfh cells, as studies in other immune cells like Tregs and B cells have shown that its deletion enhances IFN*γ* production^33; 41^. Moreover, FOXP1 is known to regulate genes involved in glycolysis pathway, such as hexokinase, phosphofructokinase, and lactate dehydrogenase, which are well-established FOXP1 targets. Thus, the increased expression of these genes under glycolysis-inhibited conditions maybe mediated by FOXP1 activity, as this transcription factor can directly bind to their promoter regions and enhance their expression ^42^.

Another important observation in this study is the disrupted fatty acid metabolism during inhibition of glycolysis in Tfh cells. Our transcriptomic data further revealed that most of the genes involved in fatty acid metabolism are suppressed during glycolysis-inhibition, as seen by downregulated levels of CPT1, OLAH, SREBF, and PPARA. It is worth noting the fact that higher OLAH have been recently linked to higher levels of IFNγ in diseases like influenza and Covid-19^43; 44^. It’ll be worthwhile to explore this axis in the Tfh cells to understand if OLAH mediated axis is also driving IFNγ production in these cells. Interestingly, this phenotype was not observed with the mouse Tfh cells, which showed elevated IL-21 production during DUSP6-mediated downregulation of glycolysis and showed no disruption of FAO during glycolysis-inhibition ^45^. However, our observation is corroborating with the previous report that showed glycolysis-dependent de novo fatty acid synthesis essential for deciding the differentiation fate of Th17 or Treg cells ^38^.

While the signalling axis for glycolysis mediated FA metabolism is not clear in these cells, the dependency of Tfh function on FA metabolism was evident even in absence of glycolysis-inhibition. FOXP1-axis seems to play a critical role in Tfh function, however it’s not clear if it is a direct link between Glycolysis and FA metabolism. There is also a possibility that fatty acid metabolism can regulate FOXP1 expression. Therefore, further investigations are necessary to establish the crosstalk of FOXP1 and fatty acid metabolism. The dependency of IL-21 on FA metabolism is so exclusive that even supplementation with acetate in glycolysis-restricted conditions has led to recovery of only IFN*γ*, but not IL-21. However, our analyses cannot clearly determine if the recovery of only IFN*γ* was due to corrected FA metabolism or due to a direct acetylation of IFN*γ* promoter region by acetate supplementation in glucose-restricted conditions^46^.

In summary, metabolic regulation of ex vivo human Tfh cells is an intricate phenomenon, mainly governed by glycolysis. However, this regulation is time-dependent, as early inhibition of glycolysis, but not in later phases, is essential to perturb Tfh cell differentiation and functions. In addition, inhibition of glycolysis also led to impaired fatty acid metabolism, suggesting a significant link between these two pathways. Certainly, glycolysis can differentially regulate Tfh cell differentiation and functions through complex regulation of different transcriptional axes. Our study clearly highlights that Tfh cell function is intricately linked to fatty acid metabolism, and this regulation is, in turn, modulated by glycolysis. This metabolic crosstalk underscores a previously underappreciated layer of control over Tfh biology, revealing how shifts in cellular energy pathways can directly influence immune outcomes. These insights not only deepen our understanding of Tfh cell biology but also offer potential avenues for targeted immunomodulation to enhance vaccine efficacy and immune regulation.

## Acknowledgements

We are thankful to the participants for generous support in this study. We thank the staff at AIIMS blood bank and Mr. Sudipta Das for technical help. This work was financially supported by the Department of Biotechnology (BT/PR25335/NER/95) and Core grants of National Institute of Immunology to NG.

## Authors Contribution

NG, conceptualized and supervised the study; PC, enrolled and collection of blood from healthy donors; SNJ, SS, sample collection; SNJ, study design and investigations; SNJ, BN, DJ, SKR, NG, formal analysis; SNJ, writing–original draft; NG, Reviewing and editing manuscript; NG, funding acquisition. All the authors edited and reviewed the final manuscript.

## Conflict of Interest

The authors declared no commercial or financial conflicts of interest.

## Notes

### Competing Interest Statement

The authors have declared no competing interest.

